# GEMsembler: cross-tool structural comparison and ensemble modeling improve metabolic model performance

**DOI:** 10.1101/2025.04.09.647174

**Authors:** Elena K. Matveishina, Bartosz J. Bartmanski, Sara Benito-Vaquerizo, Maria Zimmermann-Kogadeeva

## Abstract

Genome-scale metabolic models (GEMs) are widely used in systems biology to investigate metabolism and predict perturbation responses. Automatic GEM reconstruction tools generate GEMs with different properties and predictive capacities for the same organism. Since different models can excel at different tasks, combining them can increase metabolic network certainty and enhance model performance. Here, we introduce GEMsembler, a Python package designed to compare cross-tool GEMs, track the origin of model features, and build ensemble models containing any subset of the input models. GEMsembler provides comprehensive analysis functionality, including identification and visualisation of biosynthesis pathways, growth assessment and an agreement-based curation workflow. GEMsembler-curated ensemble models built from four *Lactiplantibacillus plantarum* and *Escherichia coli* automatically reconstructed models outperform the gold-standard models in auxotrophy and gene essentiality predictions. Optimising Gene-Protein-Reaction (GPR) combinations from ensemble models improves gene essentiality predictions, even in the manually curated gold-standard models. GEMsembler explains model performance by highlighting relevant metabolic pathways and GPRs alternatives, informing experiments to resolve model uncertainty. Thus, GEMsembler facilitates building more accurate and biologically informed metabolic models for systems biology applications.

## Introduction

Genome-scale metabolic models (GEMs) are among the fundamental tools in systems biology used to describe cellular metabolism and predict perturbation responses (Oberhardt et al., 2009; Øyås & Stelling, 2018). GEMs are reconstructed based on genome annotations and represent a metabolic network of reactions and metabolites associated with enzymes via Gene-Protein-Reaction (GPR) rules. Flux balance analysis (FBA) and its variations are used to estimate metabolic fluxes in the network under given conditions, thus predicting growth and consumption and production of metabolites, while allowing integration of different types of experimental data to constrain the model (Fang et al., 2020; O’Brien et al., 2013). The model quality is therefore crucial for accurate model predictions (Thiele & Palsson, 2010).

Whereas manual curation remains the gold-standard for production of high-quality models (Thiele & Palsson, 2010), there are various tools for the automatic reconstruction of draft GEMs that utilise different approaches and can be used as a starting point (Mendoza et al., 2019). Some tools, like gapseq (Zimmermann et al., 2021) or modelSEED (Henry et al., 2010; Seaver et al., 2021), follow a bottom-up approach by mapping enzyme genes found in the genome to the known reactions from the biochemical databases and subsequently filling the gaps to form a complete network. An alternative top-down approach is proposed in the CarveMe tool (Machado et al., 2018), which starts with a universal model from the BiGG (King et al., 2016) database and carves out unnecessary reactions based on the enzyme presence. For human gut bacteria, there is a widely used AGORA collection of semi-automatically built models (Heinken et al., 2023; Magnúsdóttir et al., 2017), which can be downloaded from the Virtual Metabolic Human database (Noronha et al., 2019). Each GEM reconstruction tool has its own advantages, and none of the tools consistently outperforms the others (Mendoza et al., 2019).

GEMs built by different tools for the same organism often use different database nomenclatures, have different structures and functional performance, making a direct comparison challenging (Fang et al., 2020). For example, modelSEED models are built using the modelSEED database (Seaver et al., 2021), gapseq relies on several integrated databases, including ModelSEED (Seaver et al., 2021) and MetaCyc (Caspi et al., 2020), while CarveMe uses the BiGG database (King et al., 2016). One approach for unifying nomenclature is offered by MetaNetX (Ganter et al., 2013; Moretti et al., 2021), an online platform that connects metabolites and reactions namespaces from different databases. However, while comparing lists of metabolites and reactions provides an overview of models’ similarities, the structural and functional differences between GEMs are not revealed. Alternatively, one can compare models based on their functional performance, such as prediction of growth, auxotrophy, or gene essentiality compared to the experimental data (Bernstein et al., 2023; Loghmani et al., 2024; Moutinho et al., 2022; Ong et al., 2020; Xavier et al., 2018). However, this approach does not reveal the differences in the network structures of models constructed with different tools.

Emerging cross-tool studies (Hari et al., 2024; Hsieh et al., 2024; Wendering & Nikoloski, 2022), show that models built with different tools can capture various aspects of metabolic behaviour, and therefore combining them in one model may improve the model performance. Several frameworks were proposed to merge GEMs built with different tools. For example, modelBorgifier (Sauls & Buescher, 2014) allows merging of two models in a semi-automated manner, while mergem (Hari et al., 2024) automatically generates a union model from several input GEMs containing metabolites and reactions from the original models. However, to date, there is no framework that can merge all model features, including genes and GPRs, track the origin of each feature in the output model, and generate fine-tuned, flexible combinations of GEMs.

To address this need, we developed GEMsembler, a python package for comparing, combining and analysing GEMs built with different tools. GEMsembler has the following unique features: i) it enables structural comparison of GEMs built with different tools; ii) it systematically assesses confidence of the metabolic network at the level of metabolites, reactions and genes; iii) it provides a comprehensive framework for assembling different combinations of the input models and assessing their predictive capacity in terms of growth, auxotrophy and gene essentiality. Further, ensemble models generated from the input GEMs can be curated using the GEMsembler functionality in a semi-automated manner. Using the two model organisms *Escherichia coli* and *Lactiplantibacillus plantarum* (formerly *Lactobacillus plantarum*) as examples, we demonstrate that GEMsembler-curated ensemble models can outperform the current gold-standard manually curated models in auxotrophy and gene essentiality predictions. Furthermore, GEMsembler framework highlights the features that explain the improved model performance, thus providing valuable information for targeted experimental validation to elucidate the knowledge gaps and uncertainties in metabolic networks.

## Results

### Generating cross-tool ensemble models with GEMsembler

GEMsembler assembles GEMs following a workflow consisting of four major steps: i) conversion of features of the input models (metabolites, reactions and genes) to the same nomenclature, ii) combination of the converted input models into one object, which we refer to as supermodel, iii) generation of ensemble models containing different combinations of the input models’ features, and iv) comparison and analysis of the ensemble models (Fig. 1A, S1).

**Figure 1.**
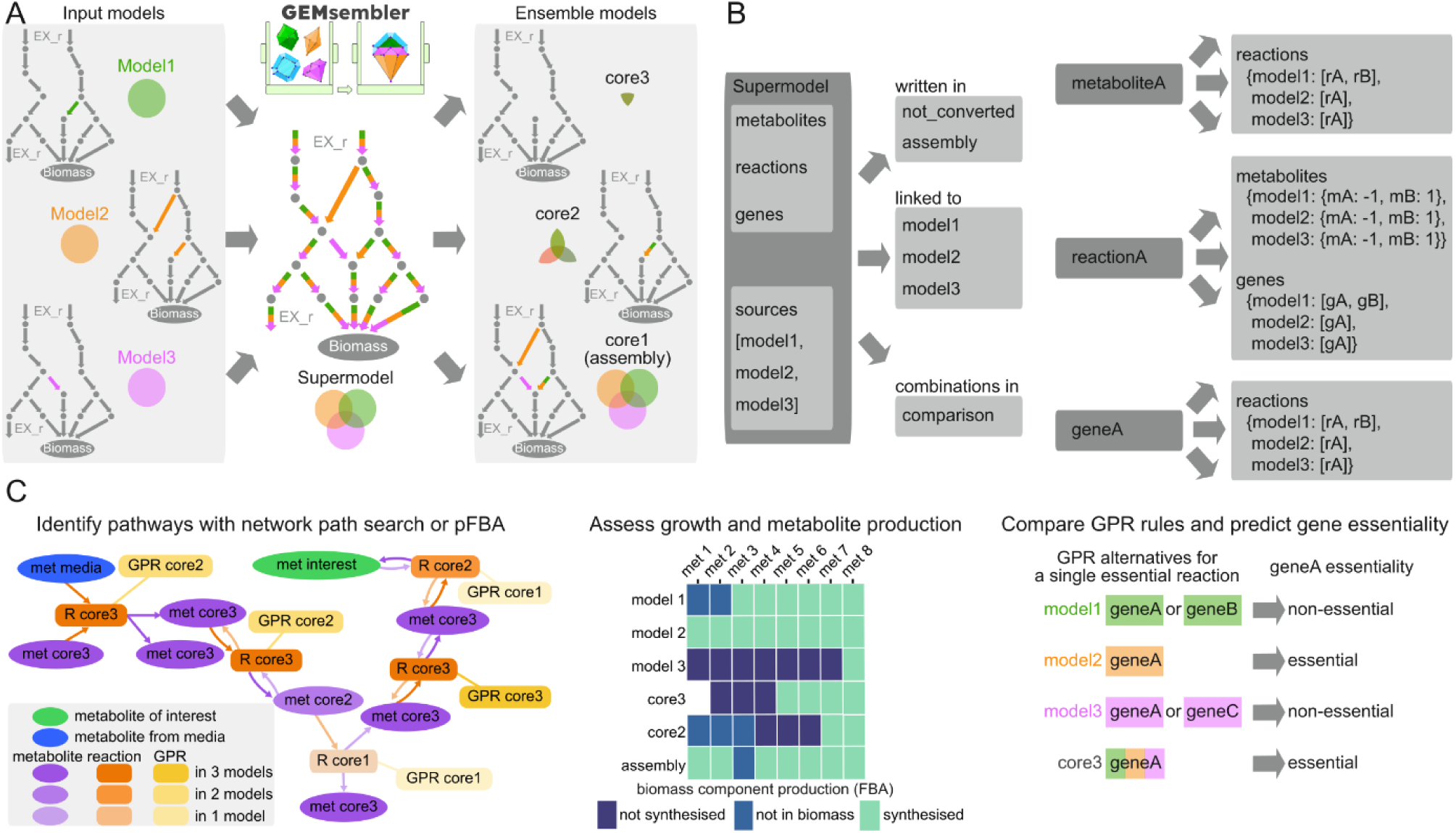
GEMsembler converts metabolic networks built with different tools to one nomenclature, combines them into supermodel, and offers diverse comparison functionality. **(A)** Schematic representation of GEMsembler workflow including construction of supermodel and generation of ensemble models with different confidence levels. **(B)** Supermodel structure resembles the structure of a COBRApy model Python class with additional information on feature conversion and their original sources. **(C)** Examples of the downstream analysis functionality, including pathway visualisation (left), growth analysis by assessment of metabolite production (center) and testing gene essentiality via GPR combination (right).

First, GEMsembler converts metabolite IDs of the input models to BiGG IDs (King et al., 2016) using different sources of information linking IDs from various databases and BiGG. Next, converted metabolites are used to convert reactions to BiGG nomenclature via reaction equations (Fig. S1) to ensure that the converted model maintains the same topology as in the original models. Finally, if genome sequences are provided along with the input models, GEMsembler converts genes from the input models to the locus tags of a genome selected as the output using BLAST (Camacho et al., 2009) (Fig. S1). GEMsembler keeps track of all the intermediate conversion steps, providing the possibility to troubleshoot in case of conversion issues.

After performing the conversion, GEMsembler assembles all converted models into one supermodel (Fig. 1A, S1). The supermodel follows the structure of the COBRApy python class (Ebrahim et al., 2013), while having additional fields to store the information about the converted features (metabolites, reactions or genes) and their source of origin (Fig. 1B, S1). Features that could not be converted are stored in a separate field (termed ‘not_converted’) in the supermodel. During the creation of the supermodel, only the union of the input models (termed ‘assembly’) is generated, which includes all features present in at least one model. All the other possible combinations of the input models, termed ‘ensemble’ models, are generated in the next workflow step. For example, we can generate ‘coreX’ ensemble models with features that are included in at least X of the input models (assembly is therefore identical to the core1 model). We define the feature confidence level as the number of input models which include this feature. The feature attributes in the ensemble models are assigned according to the same agreement principle. For example, if a reaction is unidirectional in three out of four input models and bidirectional in one model, it will be unidirectional in core4, core3 and core2 ensemble models and bidirectional in the assembly (core1) model. Gene-protein-reaction rule attributes (GPR) of reactions are compared based on the logical expressions involving genes from the original models to create new GPRs for the output ensemble models. The ensemble models are stored as a field in the supermodel object and can be extracted as separate models in the standard SBML format for downstream analysis with COBRA tools, such as flux balance analysis, gene essentiality prediction, and other functions.

### Investigating the structure and functions of the metabolic network with ensemble models

The GEMsembler supermodel consists of metabolites, reactions and genes objects, which can be examined interactively, and provides information about the original models’ agreement on the network features, which can be used to assess their confidence and identify gaps that could be experimentally validated. Due to the large number of reactions, metabolites and genes present in GEMs, it is challenging to identify subnetworks or features of particular interest for further exploration. To facilitate this analysis, we integrated additional functionality in GEMsembler (Fig. 1C) to systematically investigate the structure and functions of the metabolic network and identify subnetworks with various levels of confidence using ensemble models.

To explore the network structure, GEMsembler implements neighbourhood search to identify all reactions at a given distance from a metabolite of interest, as well as three ways of defining a pathway, for which the confidence should be assessed. First, the pathway can be predefined by the user (*e.g.*, glycolysis, tricarboxylic acid (TCA) cycle). Second, the pathway can be calculated from the network by identifying all possible ways of synthesising metabolites of interest in a given culture medium using a topology approach. In this approach, all possible paths (sequences of reactions) between the input medium components and the target metabolites are calculated by the MetQuest package (Ravikrishnan et al., 2018) integrated into GEMsembler. This topology approach generates pathways that resemble classically defined biochemistry pathways, for example, in the KEGG (Kanehisa & Goto, 2000) or MetaCyc (Caspi et al., 2020) databases. While these pathways exist in the network, it is not guaranteed that they can carry flux, as this approach does not perform flux analysis at the network scale. Therefore, the third approach to determine the biosynthesis pathways is to perform parsimonious flux balance analysis (pFBA) (Lewis et al., 2010), optimising the production of each metabolite of interest, while identifying the solution with the minimal flux. Once the pathway is defined, its confidence can be assessed by comparing the agreement scores for its metabolites, reactions and the corresponding GPR rules between the input models. This information is provided as an output table and as an interactive pathway map that can be explored visually (Fig. 1C).

To investigate the model functions, GEMsembler assesses the models’ ability to grow in a given medium by performing FBA using biomass production as the objective function. Next, GEMsembler performs FBA using the production of each component of the biomass reaction as the objective function to identify which metabolites can not be produced in case the models are not able to grow (Fig. 1C). Finally, GEMsembler incorporates GPR rules predicted by the different models and assembles their combinations. Having several GPR options per reaction can be used to guide model curation with respect to the gene essentiality prediction (Fig. 1C).

### GEMsembler enables systematic characterisation of uncertainties in GEMs

To demonstrate GEMsembler functionality, we investigated GEMs of two well-studied bacteria: *Lactiplantibacillus plantarum* (formerly *Lactobacillus plantarum*) *WCFS1* (LP), a gram-positive bacterium living in fermented foods and gastrointestinal tract, auxotrophic to multiple nutrients, and *Escherichia coli BW25113* (EC), the best studied gram-negative bacterium with a large collection of gene essentiality data. Since both of these organisms have manually curated GEMs (Monk et al., 2017; Teusink et al., 2005), we can use them as gold-standards for comparison with the outcomes of GEMsembler workflow. We set out to i) automatically generate GEMs for the two organisms with four different tools: CarveMe, gapseq, modelSEED and AGORA; ii) assemble supermodels with GEMsembler; iii) assess the agreement of metabolic reactions and GPRs by investigating different ensemble models: from the union of all models (core1) to the intersection of all models (core4 - since we are investigating four input models per bacteria); iv) compare the ensemble models to the gold-standard ones.

The majority of metabolites, reactions and genes from the four original models were successfully converted and included in the supermodels (Fig. S2, Table S1). Approximately half of the metabolites and reactions were identified only by one model, while between a quarter and a third of the genes had confidence level core1 (LP: 339 out of 1186 genes; EC: 663 out of 1952 genes). Complete agreement between all four models was observed for no more than a quarter of metabolites, reactions and genes. For each reaction with a GPR rule, we also calculated GPR agreement, which often does not correspond to the agreement score of the reaction itself. In general, the agreement between *E. coli* models is higher than between *L. plantarum* models (Fig. S2, Table S1).

To investigate agreement between different pathways, we focussed on the production of central carbon metabolites (Fig. 2A, B, Table S2), and biomass components (metabolites included in the biomass reaction) (Fig. S3, Table S3). We first identified topologically possible biosynthesis pathways in PMM5 minimal medium reported for *L. plantarum* (Wegkamp et al., 2010) and in M9 minimal medium for *E. coli*. Next, we assessed the confidence of each identified pathway by checking the confidence of reactions and GPRs in each path. We found that for most of the central carbon metabolites, the models unanimously agree on whether they are produced or not. All four *L. plantarum* models also agree on the way most of the central carbon metabolites are produced, although the GPR agreement is lower (Fig. 2A). For *E. coli,* there is absolute agreement between models for all cases except one, beta-D-glucose 6-phosphate, which includes several reactions present in three models out of four (Fig. 2B). While for all other metabolites there is core4 agreement for GPRs for almost all reactions, four reactions have GPR with core3 agreement: HEX1, EDA and ALKP involved in the synthesis paths of most of the tested metabolites, and FBA3 involved in the synthesis of D-Fructose 1,6-bisphosphate (Fig. 2B, Table S2). We also assessed the confidence of canonically defined glycolysis, pentose phosphate pathway and the TCA cycle. As expected, there is almost complete agreement in the glycolysis pathway with only some discrepancies on GPR level (Table S4, File S1, S2). Pentose phosphate pathway is also confident with disagreement in one *L. plantarum* reaction and three *E. coli* reactions out of eleven (Table S4, File S3, S4). The TCA cycle showed the most uncertainty: for *E. coli* mostly on the GPR level, but for *L. plantarum,* most of the TCA cycle is either absent or identified with large discrepancies (Table S4, File S5, S6).

**Figure 2.**
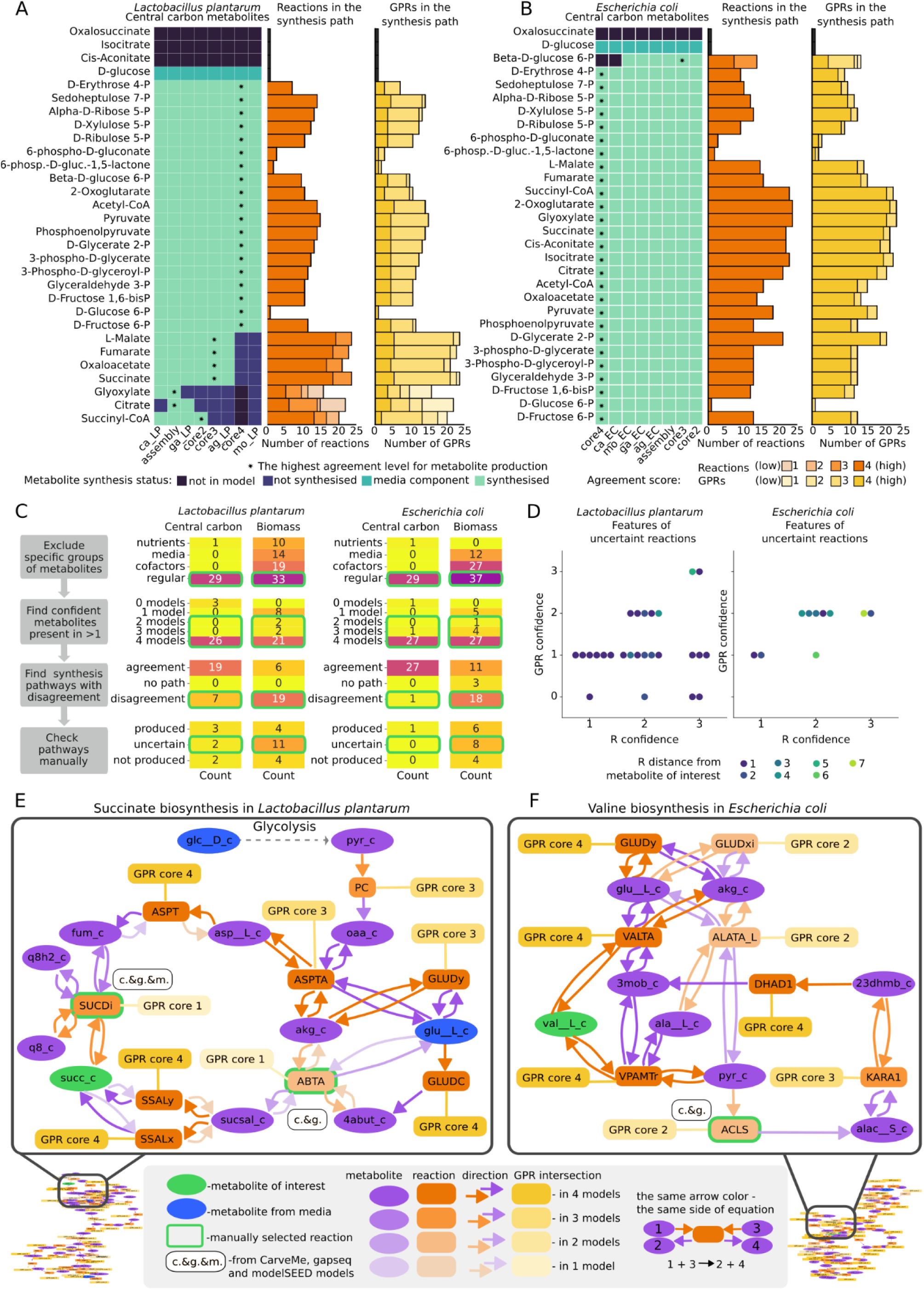
Analysis of metabolic network confidence for *L. plantarum* and *E. coli* models. **(A)** Central carbon metabolites production in the converted original models and ensemble models on the left, and agreement scores for reactions and GPRs in the corresponding most confident pathways (highlighted with *) for *L. plantarum.* **(B)** The same plots for *E. coli.* **(C)** Step-wise procedure of identification of the most uncertain pathways and reactions for further investigation (left) and the results of central carbon metabolites and biomass components examination in *L. plantarum* and *E. coli* (right). **(D)** Characteristics of the most unconfident reactions selected at the last step in C for *L. plantarum* and *E. coli* with respect to their confidence, confidence of the corresponding GPR rules and the distance from the selected metabolite of interest (which requires these reactions to be synthesised). **(E)** Example of an uncertain pathway in *L. plantarum* (succinate biosynthesis, caused by the SUCDi and ABTA reactions). **(F)** Example of an uncertain pathway in *E. coli* (valine biosynthesis, caused by the ACLS reaction).

We next performed the same topology-based analysis for the production of biomass components. Different models include different metabolites in the biomass reaction; here we decided to use their union for the complete overview (76 reactants in the union of both bacteria) (Table S5). From this analysis, we excluded metabolites that are either media components or cofactors. Production of biomass components is much less confident than that of central carbon metabolites for both organisms, but the trend that *E. coli* models have more agreement stands. For example, only 6 biomass components in the *L. plantarum* model were produced with complete agreement, while 11 biomass components had core4 agreement in *E. coli* (Fig. S3, Table S3).

Summarising the results of the topology analysis allows us to identify areas of uncertainty in the large metabolic network, which should be further investigated in a step-wise approach (Fig. 2C). First, for the metabolites of interest (28 central carbon and 76 biomass components) we separated the regular metabolites from the other types of metabolites, such as nutrients, media components (other than nutrients), and cofactors. Next, for regular metabolites, we assessed the model agreement score, and filtered out metabolites below agreement score core2. This left us with 51 metabolites for *L. plantarum* and 60 for *E. coli* model, for which we then checked production pathways. Metabolites without complete model agreement on their biosynthesis paths were then subject to manual examination (Fig. 2C, Tables S6).

In this final step, we examined the biosynthesis pathways of 7 central carbon metabolites and 19 biomass components of *L. plantarum*, as well as 1 central carbon metabolite and 18 biomass components of *E. coli* using the interactive maps generated by GEMsembler and manually grouped them into three categories: produced, not produced, and uncertain (Fig. 2C, Table S6). The pathways from the uncertain category contain reactions that are needed to produce metabolites of interest, and that have discrepancies between two or more models on the gene level, therefore they should be prioritised for further investigation. For *L. plantarum*, these include pathways for 2 central carbon metabolites, succinate and succinyl-CoA, and 11 biomass components, while for *E. coli* these include 8 biomass components (Fig. 2C, Table S6). Each of the uncertain biosynthesis pathways contains at least one reaction causing the uncertainty, with a total of 26 uncertain reactions for *L. plantarum* and 10 uncertain reactions for *E. coli* (Fig. 2D, Table S6). Most of these reactions in *L. plantarum* are confirmed by two models and have GPR included only in one model, while for *E. coli*, uncertain reactions have slightly better confidence with GPRs included in two models. Most of these uncertain reactions are in the immediate proximity to or one reaction away from the investigated metabolite, while some are up to seven reactions away in the biosynthesis pathway (Fig. 2D, Table S6).

GEMsembler provides the opportunity to explore these pathways and uncertainties visually with interactive maps and select their key elements, as we show for succinate biosynthesis in *L. plantarum* (Fig. 2E, Table S6, File S7). In this case, uncertainty is caused by two alternative paths: one through SUCDi reaction and another through ABTA reaction with GPRs provided only by one model. Another example is valine biosynthesis in *E. coli* with three reactions found only by two models: ACLS, GLUDxi and ALATA_L (Fig. 2F, Table S6, File S8). GLUDxi duplicates the function of a more confident GLUDy with the only difference in using NAD instead of NADP and, therefore, does not influence valine production. ALATA_L is required for alanine biosynthesis and therefore was considered as an uncertainty in the alanine pathway. The ACLS reaction is directly required to produce valine. Identified uncertain reactions can be either validated by existing knowledge, such as ACLS, which is confirmed by the KEGG database (Kanehisa & Goto, 2000), or serve as candidates for further experimental verification.

In this part, we demonstrated how GEMsembler helps to systematically characterise the confidence in the metabolic network and prioritise uncertain pathways and reactions for further investigation based on their confidence levels.

### Curation of GEMs with GEMsembler to reproduce growth phenotypes

The ensemble models constructed with GEMsembler can facilitate model curation to reproduce growth with the classical FBA algorithm. Since both *L. plantarum* and *E. coli* can grow in defined minimal media (Table S7), we can use this information to curate their models with GEMsembler following the agreement principle. GEMsembler growth analysis functionality uses FBA to identify biomass components which cannot be produced, therefore explaining the lack of growth under the given conditions. Preliminary growth simulations for mixed original and ensemble models demonstrated that neither of the models could grow, because not all of the biomass components could be produced (Table S8).

To curate the models, we decided to first modify the biomass reaction, and then make sure that all biomass components can be produced with FBA. We used the agreement score calculated by the GEMsembler biomass analysis function to keep biomass components included by three or more models (Table S5). Several metabolites with core1 and core2 agreement were also included, if they could be synthesised by the corresponding models (Materials and methods). In total, we included 58 and 61 components in the biomass reactions of *L. plantarum* and *E. coli*, correspondingly (Table S5).

After modifying the biomass reaction based on the agreement principle, we ran FBA for all models again to test whether they gained the ability to grow. Only the assembly model was able to grow, while each of the other models was not able to produce at least one biomass component (Fig. 3A, 3B, Table S8). Here, we used FBA instead of topology analysis to ensure that the metabolite production is relevant for the growth simulations and takes into account transport and cofactor utilisation. The number of metabolites produced with complete agreement (core4) was low in both organisms (LP: 10/58; EC: 12/61), but *E. coli* can produce more biomass components with higher agreement than *L. plantarum*. For *E. coli*, the core2 ensemble model can produce the majority of biomass precursors (Fig. 3A), while for *L. plantarum* core2 and core3 ensemble models were similar in their metabolite production capacity and could produce less than half of the target metabolites (Fig. 3B), underlining that model consistency is much lower for *L. plantarum* compared to *E. coli*. To balance the confidence, complexity and function (being able to grow) of the curated model, we chose core3 ensemble model as the basis, and added a subset of reactions from the other models to ensure that all biomass precursors can be produced.

**Figure 3.**
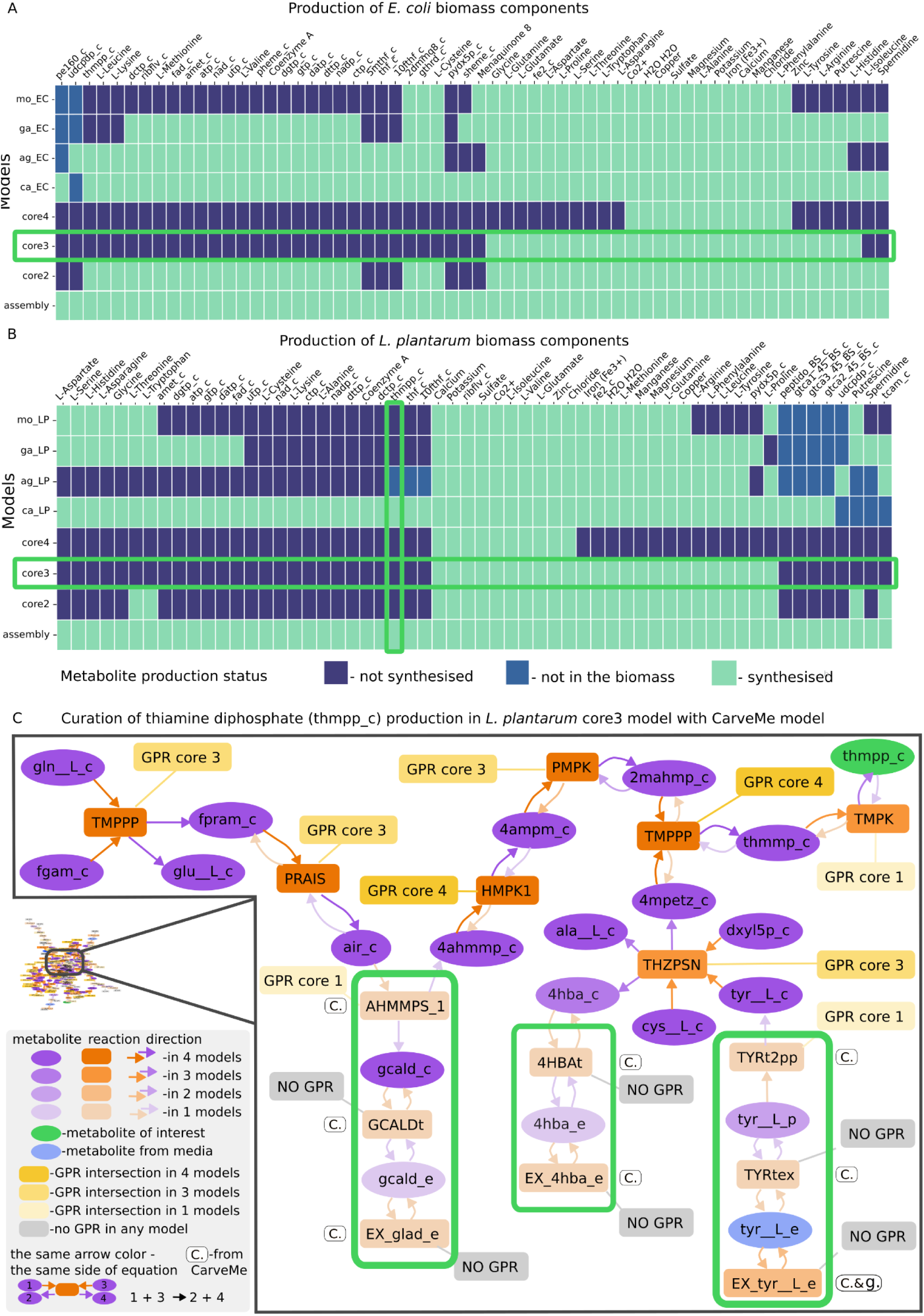
Curating *L. plantarum* and *E. coli* models with GEMsembler. **(A)** Production of unified biomass precursors for *L. plantarum* estimated with pFBA. **(B)** Production of unified biomass precursors for *E. coli estimated* with pFBA. Horizontal green boxes highlight the models selected for curation; the vertical green box highlights the metabolite for which the curated pathway is depicted in (C). **(C)** Example of curated reactions in thiamine diphosphate (tmpp_c) biosynthesis pathway of *L. plantarum*.

To determine which reactions are necessary to add for curation, we examined the interactive maps depicting pFBA-determined biosynthesis pathways for biomass components that can not be produced by core3 model, but can be produced by some other models. Reactions with less than core3 confidence level in these pathways from the other models can restore the production of the target metabolite even if they are not in its immediate proximity, but at the same time not all of them are essential for production. For example, in the path for thiamine diphosphate biosynthesis in *L. plantarum*, there are two to three unconfident reactions in three parts of the pathways that would need to be added (Fig. 3C, Table S8, File S9). One of the most challenging parts of curation is considering the highly connected biomass precursors such as ATP or NAD. In this case, sometimes adding reactions will not be enough, and reaction properties need to be changed. For example, CarveMe model was the only one able to produce ATP in *L. plantarum*, which was due to the bidirectionality of the phosphoribosylaminoimidazole carboxylase reaction (AIRCr), therefore we implemented this bidirectionality in the curated core3 model as well.

Overall, we added 72 reactions to core3 *L. plantarum* model, including 28 transport and exchange reactions, leading to the final core3 GEMsembler curated model with 639 metabolites, 729 reactions and 420 genes. We curated core3 *E. coli* model in the same way to ensure its growth on M9 minimal medium. We added 43 reactions, including 11 for transport and exchange, and the final core3 GEMsembler curated *E. coli* model consists of 943 metabolites, 1217 reactions and 644 genes. The technical quality reports generated by MEMOTE (Lieven et al., 2020) for the final core3 models and the original models confirm that GEMsembler curation solves the issue of blocked biomass precursors, that exists in all original models except CarveMe, and leads to a more realistic growth for *L. plantarum*.

In this section, we demonstrated how GEMsembler can aid model curation in a semi-automated way in order to ensure that the model can reproduce a known growth phenotype.

### GEMsembler-curated model outperforms the gold-standard *L. plantarum* model in auxotrophy prediction

After curating the core3 model of *L. plantarum* for growth in PMM5 minimal medium, which contains several amino acids and vitamins, we wanted to assess the model quality by testing whether it can predict *L. plantarum* auxotrophy to different nutrients. We compared the core3 model’s topology and performance to a previously reported curated model of *L. plantarum* iLP728 (Mendoza et al., 2019; Teusink et al., 2005), which we considered to be the gold-standard, and to the four original AGORA, CarveMe, gapseq, and modelSEED models of *L. plantarum,* converted to BiGG nomenclature with GEMsembler and automatically gap-filled with the CarveMe tool on the minimal PMM5 medium, which we considered to be the baseline.

We compared how many reactions and genes intersect between each of the models and the gold-standard iLP728 model, and calculated the ratio of each intersection to each of the model’s size (precision) or to the iLP728 size (recall) (Fig. 4A, Table S9). We also calculated the F1-score (the harmonic mean of precision and recall) as a summary metric of the reaction and gene overlap. Compared to the original input models, the core3 model recalls a similar fraction of reactions and genes corresponding to these reactions in the iLP728, while including much fewer reactions and genes not present in the iLP728 model, leading to the best F1-score for reactions (0.58) and second best F1-score for genes (0.51) (Fig. 4A, Table S9). Thus, the GEMsembler curated core3 model is more similar to the gold-standard iLP728 model of *L. plantarum* than the four original models.

**Figure 4.**
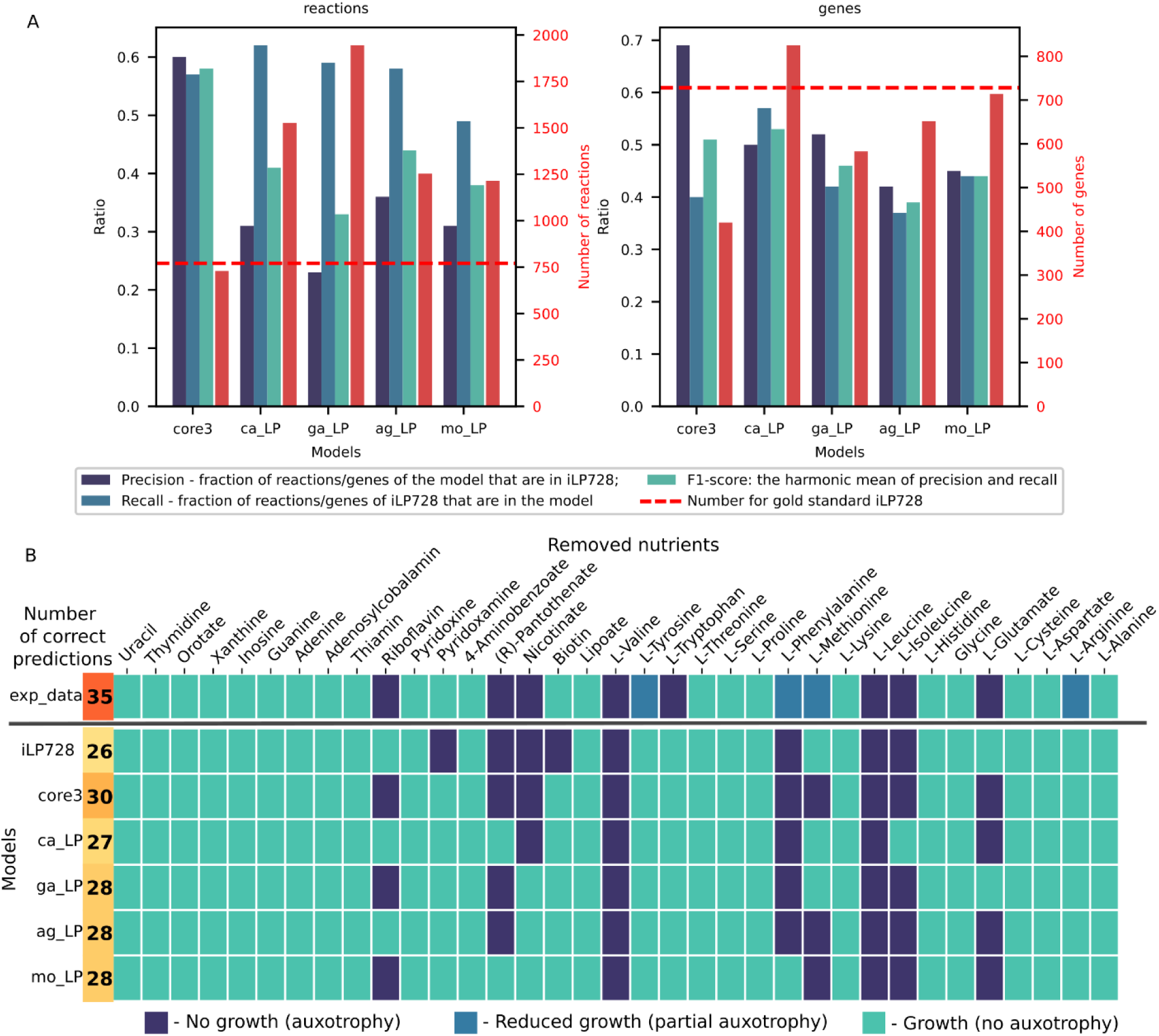
Core3 *L. plantarum* model curated with GEMsembler outperforms the other models in auxotrophy prediction. **(A)** Comparison of reactions and genes included in the core3 model, the four original models and the gold-standard iLP728 model of *L. plantarum*. **(B)** Prediction of auxotrophy to different nutrients in CDPM medium for all models compared to the experimental information. core3 - core3 model curated with GEMsembler; ag_LP - AGORA, ca_LP - CarveMe, ga_LP - gapseq, mo_LP - modelSEED models of *L. plantarum,* exp_data - experimental data.

Next, we tested the models’ ability to predict *L. plantarum* auxotrophy to different nutrients in a different medium, CDPM (Table S7), and compared the results to the reported experimental auxotrophy data classified into three categories: growth, no growth and reduced growth (Teusink et al., 2005; Wegkamp et al., 2010). We predicted growth for each model in each condition by running FBA in a modified medium where one of the tested nutrients was removed. We classified FBA prediction results into growth and no growth since there were no cases with reduced growth prediction. Our core3 model outperforms all the other models, including iLP728, in two to four growth conditions. There are no metabolites for which auxotrophy prediction is incorrect in the core3 model but correct in any other model (Fig. 4B, Table S9). Four out of five false predictions of core3 model happen in the case of experimentally observed reduced growth, while the model predicts either growth or no growth. In the case of tryptophan auxotrophy, where the core3 and all other models predict growth, the false prediction can likely be explained by the non-metabolic inhibition by other aromatic amino acids in the medium (Teusink et al., 2005). Compared to the gold-standard iLP728 model, core3 correctly predicts auxotrophy phenotype for four metabolites: growth without biotin and pyridoxamine, and absence of growth without glutamate and riboflavin (Fig. 4B, Table S9).

The GEMsembler functionality to analyse the network structure allows us to investigate the underlying reasons for improved auxotrophy predictions. In the case of biotin, the difference is that the iLP728 model considers it as a biomass component, while none of the original models and therefore neither core3 model includes it (Table S5). Although this modification of the biomass reaction leads to the correct prediction of *L. plantarum* growth without biotin supplementation, it does not explain how *L. plantarum* synthesises biotin. Since it is known that biotin is essential for fatty acid synthesis in lactic acid bacteria (Wegkamp et al., 2010), further investigation of the biotin biosynthesis pathway is needed. The opposite is observed for riboflavin, where experimentally determined auxotrophy is not predicted by iLP728, because riboflavin is not included in the biomass reaction. The core3 and all the original models include riboflavin in the biomass reaction (Table S5), which is wrongly predicted to be synthesised by CarveMe and AGORA models, while core3, gapseq and modelSEED models do not include its biosynthesis pathway. The other two prediction discrepancies between the core3 and iLP728 models can be explained by the difference in biosynthesis pathways. In case of pyridoxamine, the iLP728 model requires it to produce a biomass component pyridoxal 5’-phosphate via ALATA_Lr reaction that does not have a GPR rule (File S10). Core3 model, on the contrary, includes a different pathway to produce pyridoxal 5’-phosphate without pyridoxamine via PYDXS reaction (File S11), which is included in three original models without a GPR rule, therefore correctly predicts that pyridoxamine is not essential. Finally, glutamate auxotrophy is not predicted by iLP728 due to the presence of P5CD reaction, which enables glutamate production from proline included in the CDPM medium (File S12). This reaction does not have a GPR in iLP728, and is not included by any of the original models, therefore it is not included in core3, which leads to the correct prediction of glutamate auxotrophy.

In this section we demonstrated that the model assembly and curation pipeline of GEMsembler based on the agreement of the original models produces metabolic models that can accurately describe experimental phenotypes and even outperform the gold-standard curated model. Furthermore, the GEMsembler curated core3 model is the closest in terms of included reactions and genes to iLP728, while being the smallest of all tested models, providing a balance between model complexity and functional capacities. If additional data becomes available, it can be further curated with the help of GEMsembler, which provides candidate reactions for modification based on their confidence.

### Curating GPR rules with GEMsembler improves gene essentiality predictions in *E. coli*

Model quality depends not only on the network topology, determined by the reactions, but also on the genes included in the GPR rules. GPR rules can be tested by performing gene essentiality predictions, i.e. simulating growth while a certain gene is knocked out and the corresponding reactions cannot carry flux, and comparing the results to the experimental data. Gene essentiality predictions can vary between different input models, as the GPR rules can differ although the same input genome is used to construct the models. Since GEMsembler provides the GPR information from all the original and ensemble models, we aimed to investigate whether the space of possible GPRs can be leveraged to improve gene essentiality prediction by combining GPR rules from different models.

For our analysis, we selected the core3 GEMsembler curated *E. coli* model, and the original AGORA, CarveMe, gapseq and modelSEED models, converted by GEMsembler to BiGG nomenclature and gap-filled with the CarveMe tool on M9 medium. First, as for *L. plantarum,* we checked how similar the curated core3 model and the original models are to the latest curated *E. coli* model iML1515 (Monk et al., 2017) (Fig. 5A). For *E. coli,* the original CarveMe model is the closest one to iML1515, which is not surprising since the universal CarveMe model used for reconstruction incorporates reactions from BiGG *E. coli* models. Core3, while having the smallest number of reactions and the corresponding genes, has the highest precision for genes and the second highest precision for reactions overlapping with iML1515 (Fig. 5A, Table S10).

**Figure 5.**
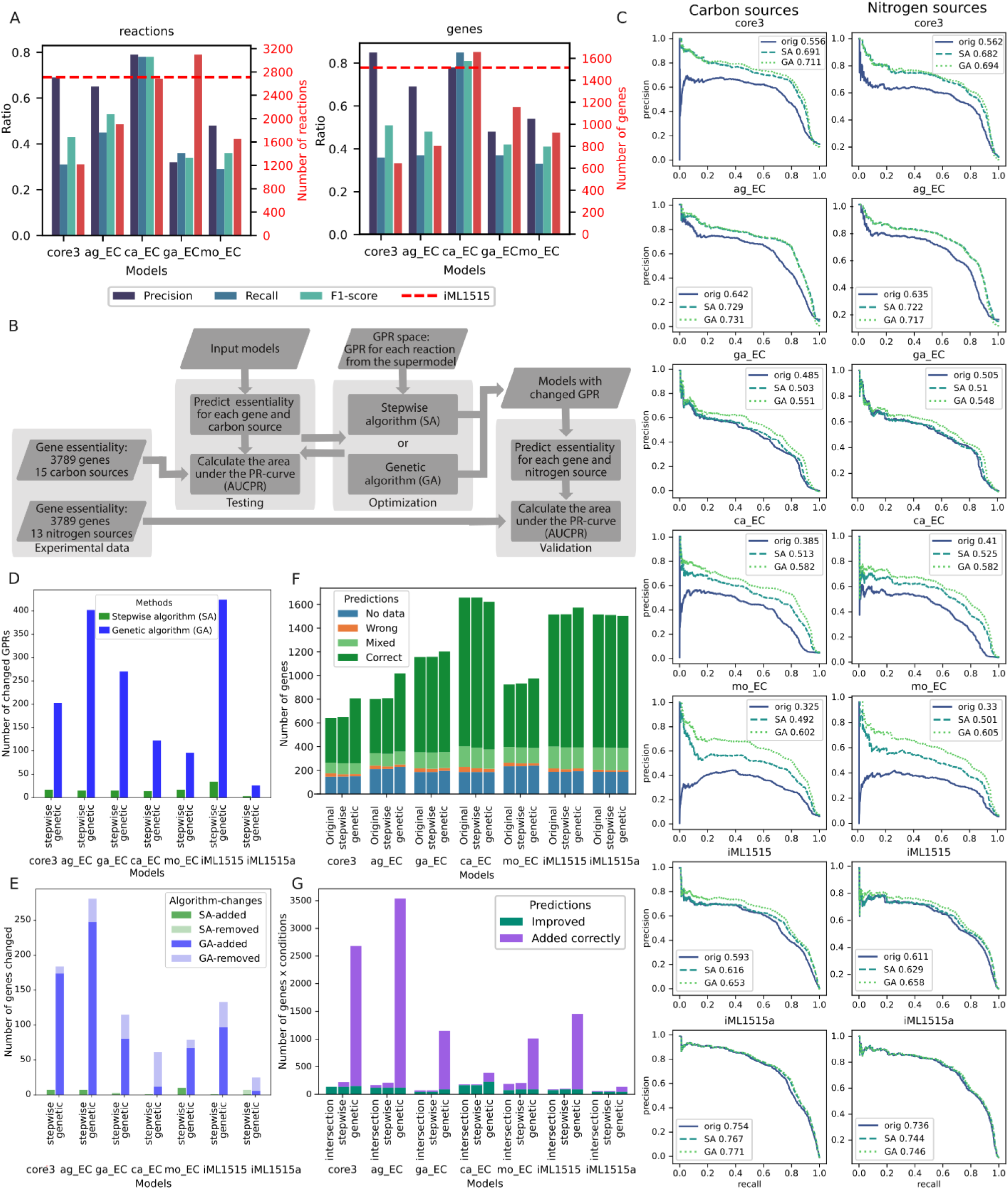
Combining GPR rules for *E. coli* models improves gene essentiality predictions for growth on different carbon and nitrogen sources. **(A)** Comparison of reactions and the corresponding genes included in the core3 model, the four original models, and the gold-standard iML1515 model of *E. coli*. **(B)** Schematic representation of the GPR modification to improve gene essentiality predictions by each model. **(C)** Precision-recall (PR) curves and their AUC for gene essentiality predictions on carbon sources (left) and nitrogen sources (right) for the original models (blue, solid) and models modified with either the combination algorithm (turquoise, dashed) or the genetic algorithm (green, dotted). **(D)** Number of changed GPRs in each model by SA and GA procedure. **(E)** Number of changed genes in each model by SA and GA procedure. **(F)** Number of genes with different prediction status for the original and SA/GA modified models using log2 fold change <-2 for fitness defect compared to the wild type to define no growth. **(G)** Number of genes for which the essentiality prediction improved using the models modified with either SA or GA, genes with a subset of correct essentiality predictions that were added to the models, and the intersection between the genes improved by SA and GA.

To assess the models’ functionality, we compared their gene essentiality predictions with experimental data on fitness defects of 3789 gene knock-out mutants in 41 minimal media, with 28 and 13 carbon and nitrogen sources, respectively (Price et al., 2018; Wetmore et al., 2015). This experimental dataset was previously used to evaluate four published curated models of *E. coli* and manually modify the latest curated model, iML1515, to further improve gene essentiality predictions and provide an adjusted model iML1515a (Bernstein et al., 2023). In our analysis we used 15 carbon sources and all 13 nitrogen sources, on which all models can grow, and calculated the area under the precision-recall curves (AUCPR) for gene-condition pairs sorted by the experimentally determined growth defect as the quality metric (Fig. 5B). The AGORA model with AUCPR=0.642 outperformed all tested models apart from the adjusted iML1515a model with AUCPR=0.754, including the standard curated iML1515 with AUCPR=0.593 (Fig. 5C). It was closely followed by the core3 GEMsembler curated model with AUCPR=0.556, while the original gapseq, CarveMe and modelSEED models perform worse with AUCPR between 0.3-0.5 (Fig. 5C).

Wrong predictions occur either due to the wrong network topology or the wrong GPR rules. We next leveraged ensemble models to address the second issue for each tested model, including iML1515 and its adjusted version. As a first approach, we implemented a stepwise combination algorithm (SA), where for each model we modified GPR rules that include wrongly predicted genes using GPR rules from the other models ordered by decreasing AUCPR, if the essentiality of the corresponding genes was predicted correctly (Materials and methods). As a second independent approach, we combined GPR rules from different models with a genetic algorithm (GA), which picks the GPR rules from all possible rules suggested by different models solely based on optimisation of the AUCPR for the carbon source conditions. After improving gene essentiality prediction for the data on different carbon sources, we tested whether the improvement was also observed for the nitrogen sources (Fig. 5B).

These two GPR modification algorithms change different numbers of GPRs in the models: while the stepwise algorithm modifies only a few dozens with minimum of three for iML1515a and maximum of 34 for iML1515, the genetic algorithm introduces hundreds of changes (Fig. 5D). These changes in GPR may also result in genes being added or removed from the model, with 1-10 genes being altered by the SA and 6-248 genes being altered by the GA (Fig. 5E, Table S10). With the SA, almost no genes were removed from the models except six genes in adjusted iML1515a model and one gene in the gapseq model. GA also tends to add new genes rather than remove existing ones except for the adjusted iML1515a and CarveMe models, which included the highest number of genes originally (Fig. 5E).

Combination of GPR rules with SA improved gene essentiality predictions for all models (Fig. 5C). The performance of the core3 model improved by 13.5% with the final AUCPR=0.691 that is higher then for the original iML1515 model. The performance of the AGORA model improved by 8.7% reaching AUCPR=0.729. Gapseq model performance was improved the least by 1.8%, while the performance of CarveMe and modelSEED models improved the most by 12.8% and 16.7% with AUCPR of 0.513 and 0.492, respectively. Even the performance of the gold-standard models could be further improved, reaching AUCPR=0.616 for iML1515 (2.3% increase) and AUCPR=0.767 for the adjusted iML1515a (1.3% increase) (Fig. 5C). While it is not surprising that models with a higher prediction quality enhance predictions of the less well performing models, our results demonstrate that model combination can also be beneficial for the opposite cases. For example, the AGORA model was improved by each of the core3, gapseq, CarveMe and modelSEED models, which have worse original prediction quality (Table S10).

Using GA to find an optimal GPR combination from the original models improves the gene essentiality predictions for all models even more than SA. The modified core3 and AGORA models reached AUCPR of 0.711 and 0.731 respectively, outperforming the modified iML1515 (Fig. 5C). Gapseq performance improved by 6.6%, while the performance enhancement for the CarveMe and modelSEED models was the largest, with improvement of 19.7% and 27.7% leading to AUCPR of 0.582 and 0.602, respectively. The performance of the gold-standard models could be improved by 6% for iML1515 and by 1.7% for the adjusted iML1515a (Fig. 5C).

Improvement in the overall prediction quality is followed by the increase in the numbers of genes with all correct or mixed predictions for all 15 carbon sources and decrease in the number of genes with the wrong predictions (Fig. 5F). Specifically 3-15 genes had wrong predictions in different original models, which got improved by either SA or GA procedure (Fig. 5G) across multiple conditions (Fig. S4). Additionally, new genes (1-8 genes for SA and 6-229 genes for GA) with correct or mixed predictions were introduced to the models (Fig. 5G, Table S10). There are no genes in any of the models that moved to the wrong prediction category due to SA or GA modifications, and only one gene was added by SA to the ModelSEED model, which had wrong predictions in all 15 carbon sources.

We investigated what genes and GPRs were introduced to the best curated model available to-date, the adjusted iML1515a, that improved its performance even further. GA and SA both improved the predictions for three genes *(b0131, b0134, b0778)* that were in the model, and introduced one gene *(b1593)* with correct predictions. Each of the improved three genes corresponds to a single reaction in the adjusted iML1515a (ASP1DC, MOHMT, DBTS respectively). These genes were originally predicted as essential, but became non-essential after the GPR modification in agreement with their non-essentiality across all experimental conditions. They became non-essential, because the corresponding GPR rules that included only one gene were changed to a GPR rule including two or three isoenzymes. Thus, GEMsembler suggests candidates for taking over the function of these genes. For example, the GPR rule of dethiobiotin synthase reaction (DBTS) was changed from (*b0778*) to (*b0778* or *b1593*). Indeed, *b1593* is annotated in Ensembl (Howe et al., 2020) as putative dethiobiotin synthetase, and itself has the correct prediction of being non-essential corresponding to the experimentally determined non-essentiality.

Since it is important to test whether the model improvement can be generalised, we performed gene essentiality predictions for the same mutants grown on different nitrogen sources (Fig. 5C). For the input models and the modified models (with GPR changed based on the carbon sources data), the improvement trends as well as the overall prediction quality was reproduced in the nitrogen sources dataset (Fig. 5C, Table S10).

GEMsembler combines and compares GEMs built with different tools, creates ensemble models and offers a diverse functionality for their analysis, balancing manual and automatic approaches. Its applications are particularly useful for assessing metabolic network confidence and curating of the models on both reaction and gene levels, thereby, increasing model performance while reducing the time invested in the standard manual curation process.

## Discussion

GEMsembler offers a comprehensive and flexible solution for comparing GEMs built using different tools and generating their ensembles. Comparison across tools is essential, since the choice of reconstruction tool can affect the model structure and predictions even more than the input genome. Indeed, it was recently shown for environmental bacterial communities, that models built with the same tool were more similar to each other in predicting exchanged metabolites than models built for the same type of community (Hsieh et al., 2024). GEMsembler reveals the variability introduced by different tools, and provides a functionality to navigate in large metabolic networks and build model ensembles based on the model agreement, potentially reducing tool-dependent biases. We demonstrate that building ensemble models is beneficial, because models can improve each other’s performance, for example in auxotrophy and gene essentiality predictions. Furthermore, comparing alternative reactions and GPR rules between different models reveals uncertainties and gaps in our knowledge of the organism’s metabolic pathways, and helps to prioritise and design experiments to elucidate these gaps. Finally, we propose a systematic and transparent workflow for curating ensemble models, demonstrating that the GEMsembler-curated models have similar quality or even outperform the gold-standard curated models of *L. plantarum* and *E. coli*. In this way, GEMsembler can streamline the time-intensive process of model construction and curation while ensuring high-quality models.

GEMsembler also has its limitations, the first being its dependency on BiGG as the primary database, which has fewer metabolites and reactions than for example the modelSEED database. We chose BiGG because it is model-oriented, widely used in the research community and has self-explanatory IDs. The flexibility of GEMsembler, however, would allow us to change the primary database, if there is a need in the modelling community. The second limitation is that GEMsembler loses unconverted metabolites, reactions and genes in the default mode of analysis. We addressed this issue by introducing a mixed option for supermodel generation, which includes non-converted network elements that can be added if they turn out to be important for the analysis. Additionally, while GEMsembler currently supports conversion from six model types, the existing model conversion classes can be adapted by the user to new model types.

One of the challenges of ensemble model building that GEMsembler attempts to solve is the ambiguous definition of combinations of specific model attributes, such as GPR rules, reaction boundaries or directionality. For instance, how is an intersection of two GPR rules defined, if in one model the rule is “(A and B) or C” and in another one “(A and D) or E”? On the one hand, it can be A, but on the other hand, it can be empty. We implemented the former option as the default, but included a “and_as_solid” parameter to all functions that include a GPR combination, which can be changed to “True” to choose the latter option. Another combination-related challenge is the numerous ways to combine different models. In this work, we used a rigid approach, which kept model features and their attributes on which a fixed number of models agree. It can result, for example, in cases when a reaction is present because three models agree on it, but its GPR rule is not retained because there is no overlap between any three models’ GPR rules. In the future, we plan to implement a more flexible way to combine feature attributes, such as retaining a GPR rule that corresponds to the highest non-empty model agreement for each reaction. Further GEMsembler improvements could include refining the merging of the duplicated reactions, adding metabolite formulas and annotation to the output models, and checking the mass/charge balance of the combined reactions.

While here we demonstrated that GEMsembler can compare, combine and aid in the curation of models for single bacterial species, its functionality is not limited to these examples. Alternative pathways identification between different models can inform engineering strategies for strain design to increase production of metabolites of interest or optimise growth (King et al., 2015). Further applications can include comparing models of closely related species or strains to identify their metabolic differences (Bosi et al., 2016; Hari et al., 2024; Monk et al., 2013; Neal et al., 2024; Seif et al., 2018). Another application could be comparing different organisms with a similar metabolic phenotype in order to find common pathways that could be responsible for that (Oberhardt et al., 2011; O’Brien et al., 2015). Furthermore, the use cases of GEMsembler can be expanded to microbial communities (Henry et al., 2016; Machado et al., 2018; Schäfer et al., 2023). An assembly model of the entire community can provide insights on the pathways that are overlapping or complementary between different community members, suggesting potential interspecies interactions and guiding experimental design (Bauer et al., 2017; Weiss et al., 2022).

Taken together, we believe that GEMsembler’s systematic approach to ensemble modelling and ease of use will benefit researchers at any experience level interested in building, analysing and curating GEMs of their species of interest. At the same time, GEMsembler’s functionality can be generalised and integrated into other computational pipelines addressing various aspects of bacterial metabolism. We therefore hope that GEMsembler will be adopted by a broad scientific community working in the area of systems and computational biology, and contribute to building more comprehensive, concise and biologically informed metabolic models.

## Materials and Methods

### GEMsembler package overview and source code availability

GEMsembler package workflow consists of four main steps: 1) input model conversion; 2) supermodel assembly; 3) ensemble model generation; 4) assessment of model agreement and functional analysis. As input, GEMsembler requires model files in SBML format with the information on which tool was used to generate the model. Currently, GEMsembler supports input models built with either of the four tools: CarveMe (Machado et al., 2018), gapseq (Zimmermann et al., 2021), MetaNetX (Moretti et al., 2021) and modelSEED (Henry et al., 2010), or downloaded from either of the two databases: AGORA (Heinken et al., 2023) and BiGG (King et al., 2016). Models from other sources can be implemented by adding a custom model type. Optional input to GEMsmebler alongside with the models are the bacterial genomes used to generate each of the input models, as well as the genome file or NCBI assembly ID (for automatic file download), which should be used for the final conversion of GPR rules between the models. As output, GEMsembler produces the supermodel that is stored in JSON format, as well as ensemble models in the standard SBML format readable by COBRApy (Ebrahim et al., 2013) or alternative COBRA and model analysis packages (Heirendt et al., 2019; Klamt et al., 2007; von Kamp et al., 2017). At the model analysis step, GEMsembler requires the input list with the growth medium components, and the metabolites or pathways of interest, and produces plots, tables and interactive pathway maps in HTML format. GEMsembler is developed in Python and requires standard Python libraries, as well as BLAST (Camacho et al., 2009), that is needed for gene conversion, and the MetQuest package for topological network analysis (Ravikrishnan et al., 2018). Out of these dependencies, only BLAST needs to be installed separately by the user. GEMsembler source code, tutorials and example notebooks are freely available at https://github.com/zimmmermann-kogadeeva-group/GEMsembler.

### Model conversion step

In the first step, to convert metabolite IDs of the input models to BiGG IDs (King et al., 2016) GEMsembler uses and prioritises several sources of crosslink information. We checked for the BiGG IDs provided in the metabolite/reaction annotation field in the input model, as well as crosslink information from the original database of the input model (ModelSEED (Seaver et al., 2021), BiGG (King et al., 2016)). If the IDs provided in the model annotation and the original database overlap, their intersection becomes the first priority for ID mapping. If there are discrepancies between the two, the IDs provided in the model annotation field are considered as the second priority, and if it is empty, the original database crosslinks are considered as the third priority. If there are no conversion results from the model annotation or the corresponding database, the fourth priority is to use some additional conversion information, such as MetaNetX (Moretti et al., 2021) with crosslinks between many databases. If still no BiGG ID is found, the fifth priority is to change the ID according to some pattern (for example, replace one last underscore with two underscores in AGORA IDs). The last and sixth priority is to check whether the ID itself without any changes can be found in the list of BiGG IDs. For each of the currently supported model types, a separate conversion step is implemented. For a custom model type, the user needs to adapt the conversion step according to the model nomenclature. The conversion results between different databases are often ambiguous. One ID in the original database can correspond to several BiGG IDs (one-to-n), several different IDs can be converted to one BiGG (n-to-one), and even several input IDs can have links to the same set of several BiGG IDs (n-to-n). For different input models that use the same biochemical database nomenclature, such as gapseq and ModelSEED models that both use ModelSEED database, checking for conversion consistency between models allows eliminating some ambiguous results. For the main track of further steps, GEMsembler selects metabolites to ensure unique one-to-one conversion, and those that do not fulfil that requirement are considered separately.

After the metabolites are converted, the reactions are converted based on their equations. First, only uniquely converted metabolites are used to generate reaction equations in the BiGG nomenclature. For metabolites that were not converted uniquely, for example, one-to-many, GEMsembler suggests different possible conversion results to compose reaction equations with the corresponding metabolites. If one of the conversion options for a particular metabolite leads to the reaction equations present in the BiGG database, that conversion is used as a conversion result for that metabolite and its reactions. Additionally, for models that do not include a periplasmic compartment, this compartment can be introduced if there are reactions for which changing a metabolite compartment from cytosolic or extracellular to periplasmic leads to correct equations present in the BiGG database. Additionally, GEMsembler merges reactions with duplicated reaction equations in the models and in the BiGG database, keeping those that are used in the most number of models.

If provided with the input genomes used to generate the corresponding original models, and either an NCBI assembly ID or a genome file in fasta format to be used for the output models, GEMsembler converts genes included in the input models to the output genome sequences. In case an NCBI assembly ID is provided, the assembly is downloaded automatically according to NCBI ID with ncbi_genome_download package (Blin, 2023) and locus tags from the assembly are used as gene IDs, otherwise the model genes are converted to the gene IDs provided by the user in the custom fasta file. Changes introduced by different tools to the gene IDs (like addition of a dot or replacement of an underscore), or gene coordinates instead of IDs used by gapseq, are taken into account during the gene conversion process, which can be different for different types of models. The final conversion is made using BLAST sequence alignment (Camacho et al., 2009).

### The supermodel structure

Information from all the input models is stored in a Python structure called supermodel. Supermodel consists of three main classes: metabolites, reactions and genes, each of which has the following main fields: “assembly”, “comparison” and “not_converted”. The field “assembly” unites the information on individual metabolites, reactions or genes from all the input models. These main classes also have fields for each individual original model for direct access. Not converted metabolites, reactions or genes are stored in the corresponding “not_converted” field. If the user does not want to lose non-converted elements of the models, it is possible to mix them with the converted ones in the “assembly” using the original non-converted IDs from the input models. Each individual metabolite, reaction or gene has similar attributes as in the COBRApy python class structure, such as “name” or “reactions” for a metabolite, “reactions” for a gene, “metabolites”, “genes”, “gene_reaction_rule”, “lower_bound”, “upper_bound” for a reaction, but inside of each attribute, separate fields are added for each individual input model and their union. These fields allow direct comparison of the values of each attribute in each of the input models and their union. Supermodel offers various built-in comparison functionalities, for instance, calculating the features on which at least or exactly X input models agree with corresponding “at_least_in” and “exactly_in” supermodel methods. Another supermodel method called “present” allows to find features that are included by a certain list of input models but are excluded by others using logical expressions. The results of all performed comparisons are stored in the “comparison” attribute, which is empty at the beginning. Afterwards, it is possible to extract any ensemble model, in standard SBML format, as well as add or remove specific reactions the user is interested in by specifying their IDs (if a reaction is added, all its attributes are added from the assembly level).

### Analysis of the network topology and function

Once the supermodel is created and the ensemble models are built, GEMsembler can be used to perform network topology and functional analysis. For the topology-determined pathway search, we adapted the MetQuest package (Ravikrishnan et al., 2018). It needs a path to the XML model file as input, formulation of the growth medium with nutritional sources as configuration file, and some other optional parameters, for example, a list of metabolites of interest to extract corresponding pathways separately. MetQuest searches for all possible paths below a certain length between the medium components and all metabolites that can be reached using only already available compounds. For an input network with over a thousand metabolites and reactions such calculations could require > 16 GB of RAM (we recommend allocating at least 32 GB of RAM). Therefore, this path search is implemented as a separate “pathsfinding” module, which enables the user to run it on a high-performance cluster, if available. In this work, we ran ‘pathsfinding’ module using the EMBL Heidelberg HPC cluster (European Molecular Biology Laboratory et al., 2020). As output, “pathsfinding” module generates a large dictionary with all possible biosynthesis pathways under the certain length in the model in the HDF5 “metquest.h5” file, and a “shortest_paths.pkl” file with a smaller dictionary with three shortest paths (by default) for each metabolite in the model, or for metabolites of interest, if specified. Such topological “pathsfinding” outputs for several original and ensemble models can be analysed and summarised together with the “run_metquest_results_analysis” GEMsembler function.

To perform functional analysis of the networks, the FBA and pFBA functions from the COBRApy package are used. Specifically, the culture medium has to be provided to simulate growth and production of all components of the biomass reaction or of metabolites of interest provided by the user. Production of a certain metabolite is considered when the flux through the demand reaction of the specified metabolite, introduced as an objective function for simulations, is higher than 0.001 mmol g_DW_^-1^ h ^-1^. Reactions which carry flux higher than 0.001 mmol g_DW_ ^-1^ h^-1^ in the corresponding pFBA results define a biosynthesis pathway for this metabolite. This biosynthesis analysis, together with further summary and analysis for a dictionary of models is implemented in the “run_growth_full_flux_analysis” GEMsembler function.

Another type of network analysis implemented in GEMsembler with the “get_met_neighborhood” function, calculates and visualises the neighbourhood within a number of reactions, which are selected by the user, starting from a given metabolite. This functionality is also used to calculate reactions’ distance from the biosynthesis product of the corresponding pathway.

As the output, GEMsember functions “run_metquest_results_analysis” and “run_growth_full_flux_analysis” generate summary production and pathway agreement plots/tables for metabolites of interest in all models. These functions also generate sets of corresponding tables that contain identified paths, as well as interactive network maps built with networkx (version 3.3) (Newman, 2003) and pyvis (version 0.3.2) (Perrone et al., 2020) libraries.

### Input models reconstruction, supermodel and ensemble models generation for *E. coli* and *L. plantarum*

For the use case analysis, the draft models of *L. plantarum WCFS1* (LP) and *E. coli BW25113* (EC) were reconstructed with CarveMe and gapseq command-line tools and modelSEED web-server. We used protein sequences for CarveMe and modelSEED and nucleotide sequences for gapseq from the *L. plantarum* assembly GCF_000203855.3, and *E. coli* sequence files from the KEIO collection https://fit.genomics.lbl.gov/cgi-bin/org.cgi?orgId=Keio. We used gram-specific templates for the models and default gap-filling without specifying any media. We also downloaded models for the species from the AGORA2 collection (Heinken et al., 2023) https://www.vmh.life/files/reconstructions/AGORA2/version2.01/sbml_files/individual_reconstructions/. As genomes were not available for the AGORA2 collection, we used the genome from AGORA1 (https://www.vmh.life/files/reconstructions/AGORA/genomes/AGORA-Genomes.zip) for *L. plantarum,* and the genome from assembly GCF_000750555.1 for *E. coli*, as it fits to the AGORA2 model in terms of gene IDs, after selecting for locus tags (https://ftp.ncbi.nlm.nih.gov/genomes/all/GCF/000/750/555/GCF_000750555.1_ASM75055 v1/GCF_000750555.1_ASM75055v1_cds_from_genomic.fna.gz).

First, for both organisms, the input models were converted with the “GatheredModels” class and its “run” method. Next, two supermodels were assembled with the “assemble_supermodel” method of the “GatheredModels” class: one is the default with “do_mix_conv_notconv” set to False, and the other one is mixed with “do_mix_conv_notconv” set to True parameter. Finally, in both cases ensemble models were created with the “get_all_confident_levels” method of supermodels and the “get_models_with_all_confidence_levels” function. These models were used further for the topological pathway analysis as well as for the growth and pFBA pathway analysis.

As a baseline for model comparison, we took four original AGORA, CarveMe, gapseq, and modelSEED models of *L. plantarum* or *E. coli,* converted to BiGG nomenclature with GEMsembler mixed approach without adding transport reactions, and automatically gap-filled on the minimal PMM5 media with CarveMe tool gap-filling command. The biomass components for *E. coli* were taken from the “assembly” field, while for *L. plantarum* the biomass components were taken from each of the original models, and then modified according to the curation procedure below. The reason for the *L. plantarum* difference is that, when using the “assembly” biomass reaction, gap-filling of the original models was not feasible.

### Model curation

The model curation consists of two major steps: i) curation of the biomass reaction ii) curation of the biosynthesis pathways of the biomass components in a given growth medium. To decide which components of the biomass reaction to keep, a preliminary growth analysis and biomass component production analysis was performed using the original models converted to BiGG nomenclature with GEMsembler default and mixed approach. To decide which biomass components to include, we used the agreement score calculated by the GEMsembler “biomass” function to keep biomass components included by three or more models (Table S5). Several metabolites were exceptions to this rule. One of them is acyl carrier protein (ACP_c), which was included by three out of four models, but none of these models could produce it in the preliminary analysis (Table S8), therefore we decided to exclude it. For *E. coli*, we removed vitamin B12 (adenosylcobalamin adocbl_c), core oligosaccharide lipid Al (colipa_c), and phosphatidylethanolamine (dioctadecanoyl, n-C18:0, pe180_c), which were included by three out of four models but not produced by either of them (Table S5, S8). We also considered metabolites with agreement by only one or two models, taking into account whether these metabolites can be produced by the corresponding models (Table S5, S8). We kept several low-confidence metabolites, such as lipids, because they are essential for biomass production, despite the large disagreements between the models. Two lipid metabolites and siroheme for *L. plantarum,* as well as one lipid for *E. coli* were removed because their biosynthesis pathways were long, non-linear and present only in one model, therefore less likely to be present (Table S5, S8). Note that the biomass reaction selection process in GEMsembler is primarily based on the presence of the biomass components in each of the input models and whether production is achieved. Therefore, experimental validation would be necessary for the user to assess the list of biomass components and their corresponding coefficients in order to obtain a normalised and more accurate biomass reaction for the species of interest.

*L. plantarum* and *E. coli* core3 models were curated to produce all biomass precursors with the following procedure. First, for each biomass precursor which could not be produced by the core3 model, we checked the biosynthesis pathway maps from the core2 model or the original models that can produce the target metabolite. Then, we manually identified a small set of reactions missing in core3 that can restore the metabolite production, and we added these reactions. If according to the pathway map for production of one biomass component, another one needs to be produced, the biosynthesis of the latter was curated first. Transport reactions for common metabolites with agreement in less than three models identified during pathway maps exploration were added as well. Finally, if after adding all reactions from the functional biosynthesis pathway, the curated core3 model was not able to produce the metabolite, we compared reaction boundaries from the original models. For example, for ATP biosynthesis in *L. plantarum,* we compared reaction boundaries between core3 model and CarveMe model, which could produce ATP, and found a discrepancy in phosphoribosylaminoimidazole carboxylase (AIRCr) reaction. Changing the reaction boundaries of AIRCr reaction to bidirectional restored ATP production in the curated core3 model.

MEMOTE quality reports (Lieven et al., 2020) were generated with the command “memote report snapshot“. The reports for four original input, four original output and GEMsembler-curated models per species are available in the git repository https://git.embl.de/grp-zimmermann-kogadeeva/GEMsembler_paper.

### Comparison with the gold-standard models

To assess the overall similarity with the gold-standard models for both *L. plantarum* and *E. coli*, we calculated how much each of the assessed models resembles the gold-standard one (recall) and how much the assessed models are confirmed by the gold-standard one (precision). For reaction level similarity, we took all reactions from the gold-standard model and each of the assessed ones, and calculated the ratio of the number of intersecting reaction IDs to either the number of all reactions in the gold-standard model (recall) or the number of all reactions in the assessed model (precision). For gene level similarity, we took a similar approach, however, we only compared genes if they were linked to the same reaction in the compared models (both GPRs from the compared models should contain the gene, but do not have to be exactly the same). Therefore, genes that are included by both compared models but correspond to different reactions were not included in the precision and recall calculation.

### Auxotrophy prediction

For auxotrophy prediction in *L. plantarum*, we classified experimental growth in the CDPM media lacking each of the 35 components tested in (Wegkamp et al., 2010) into three groups: growth (OD_600_ >= 1), no growth (OD_600_ <= 0.1) and reduced growth (OD_600_ between 0.1 and 1) (Figure 4B). For three metabolites, the growth was modified according to the information provided in a previously published auxotrophy experiment (Teusink et al., 2005). Specifically, the growth without isoleucine was set to zero (OD_600_ of 0 instead of OD_600_ 0.2), because the authors reported that *L. plantarum* cannot produce isoleucine, but growth was observed due to trace elements remaining in the medium (Teusink et al., 2005). Growth without phenylalanine and tyrosine was set to reduced growth category (OD_600_ = 0.2 instead of OD_600_ = 0.1), because the authors reported that growth was noticeable in these conditions (Teusink et al., 2005).

To predict auxotrophy with the modelling framework and compare to the experimental results, FBA was run while optimising for biomass production as the objective function for each model in each condition where one of the tested nutrients was removed from the CDPM medium. The FBA prediction results were classified into growth (growth rate >= 1 h^-1^ (absolute threshold) or growth rate >= 0.85 of the maximum growth rate calculated in the unmodified CDPM medium of the corresponding model (relative threshold)), and no growth (growth rate <= 0.001 h^-1^ or rate <= 0.15 of the initial maximum growth rate in the unmodified CDPM medium). There were no cases with reduced growth prediction for any of the tested models.

### Gene essentiality prediction

To test the models’ ability to predict gene essentiality, we used an experimental dataset with the KEIO collection of gene knockout mutants in *E. coli BW25113* https://fit.genomics.lbl.gov/cgi-bin/org.cgi?orgId=Keio (Price et al., 2018; Wetmore et al., 2015), grown on minimal media including different carbon and nitrogen sources. This dataset was previously compiled to evaluate four published curated models of *E. coli* and manually modify the latest curated model iML1515, resulting in the adjusted iML1515a model (Bernstein et al., 2023). For each gene and each condition, a fitness defect measure defined as the log2 fold change of growth compared to the wild type in that condition was available for 3789 genes in 28 carbon sources and 13 nitrogen sources. We followed the same gene essentiality prediction pipeline, predicting gene essentiality with COBRApy function “single_gene_deletion” for each of the tested models (four original models, curated core3 model, iML1515 and the adjusted iML1515a). We defined a gene as essential if the predicted growth rate was less than 0.001 h^-1^, and nonessential otherwise. We then sorted the genes according to the experimental fitness defect in increasing order, and calculated precision-recall curves and their Area Under the Curve (AUCPR) with Scikit-learn (version 1.5.1) Python library. Note that for each model, we could calculate essentiality for a different number of genes, depending on how many genes from the experimental dataset were included in the model. To make the essentiality prediction comparison more fair between the models, we used 15 out of 28 available carbon sources in the dataset and 13 nitrogen sources on which all models could grow. For the experimental gene essentiality threshold used to calculate the number of improved predictions, we used a fitness defect of log2 fold change equal to -2, as used in the previous study (Bernstein et al., 2023).

### Improving gene essentiality prediction

To improve gene essentiality predictions for each of the tested models, we modified the GPR rules in each model following two approaches. In the first approach, we designed a stepwise GPR rule combination algorithm (SA) to use GPR rules from the other models for modification of the target model. The first step is to select the reactions for which GPR contains a gene with the wrong essentiality prediction in the target model. The next step is to modify the GPR rule by changing it to the GPR rule from a different model, in which the gene essentiality prediction was correct for that gene. If the GPR modification improved the essentiality prediction of the selected gene and did not affect the prediction for the correctly predicted genes by the target model, the GPR was kept. Otherwise, a GPR from a different model for the same reaction was tested. The models which were providing an alternative GPR were tested in the order of decreasing AUCPR.

As a second approach, we implemented a genetic algorithm (GA), which aims to find a GPR rule combination in the solution space that optimises the AUCPR for gene essentiality prediction of the target model. The reactions of the input model are considered as genes in the chromosome, in the context of the genetic algorithm, if they have different GPR variants in some of the original or core3 models. Potential sources of GPR information were encoded as integers (0 for input model itself, 1 for core3, 2 for ag_EC, 3 for ga_EC, 4 for ca_EC and 5 for mo_EC). Selecting one number (one GPR source) per reaction leads to a solution vector and combining numbers for sources with different GPRs for all reactions forms the solution space for the algorithm. We search for a solution that leads to the highest AUCPR with PyGAD (version 3.4.0) python package with the following parameters: “num_generations”: 50, “num_parents_mating”: 40, “sol_per_pop”: 200, “parent_selection_type”: “tournament”, “K_tournament”: 20, “keep_elitism”: 5, “crossover_type”: “two_points”, “mutation_type”: “random”, “mutation_by_replacement”: true, “mutation_probability”: 0.05, “random_seed”: 42, “parallel_processing”: 100. The genetic algorithm optimization was executed on the EMBL Heidelberg HPC cluster (European Molecular Biology Laboratory et al., 2020). We also want to note that we encountered numerical accuracy issues when using the glpk solver package when running the “single_gene_deletion” function from the COBRApy package, which we resolved by using cplex solver instead. To reduce the number of randomly introduced GPR changes, we intersected solutions from different generations, introducing only changes on which the last N generations agree. We started from the last one (N=50), and tested the intersections of the GPRs with the previous generations in terms of the AUCPR, adding one previous generation at a time. The procedure was stopped when the resulting AUCPR decreased below the AUCPR of the 50^th^ generation rounded to the smaller value with second decimal precision.

## Supporting information

Content readme for supplementary tables and files on zenodo

## Data and code availability

GEMsembler source code, tutorials and example notebooks are freely available at https://github.com/zimmmermann-kogadeeva-group/GEMsembler.

Code and data required to reproduce results from the study are available at https://git.embl.de/grp-zimmermann-kogadeeva/GEMsembler_paper.

Data and tables used in the study are available on Zenodo: https://doi.org/10.5281/zenodo.15130953.

## Author contributions

Elena K Matveishina: Conceptualization; data curation; software; methodology; formal analysis; validation; visualization; investigation; writing – original draft; writing – review and editing.

Bartosz J. Bartmanski: Conceptualization; Data curation; software; formal analysis; validation; visualization; writing – review and editing.

Sara Benito-Vaquerizo: Conceptualization; Validation; writing – review and editing.

Maria Zimmermann-Kogadeeva: Conceptualization; supervision; validation; visualization; resources; project administration; funding acquisition; writing – review and editing.

## Statement on using AI tools

In the preparation of this research paper, we utilised generative AI tools (ChatGPT 4o and perplexity.ai) to enhance the clarity and coherence of our writing. We carefully reviewed and edited the AI-generated recommendations to maintain the integrity and authenticity of our work. Generative AI-tools were not used to produce new content, but only to rephrase human-written text to improve its clarity. ChatGPT 4o was also used to suggest code implementation to generate figures, which was subsequently edited by the authors.

## Acknowledgements

We would like to thank members of the Zimmermann-Kogadeeva lab for helpful discussions and feedback. We would like to acknowledge Gleb E. Gavrish for his help in cross-platform integration of GEMsembler, Artemiy Golden for his input on supermodel creation algorithm, Jean-Karim Hériché and Renato Alves for advice on the implementation of the genetic algorithm, and Santhust Kumar for advice on pathways identification. The work was supported by: the European Molecular Biology Laboratory; the EMBL International PhD Programme (E.K.M.); the European Research Council (ERC) (MetaboGutModel-101117769), and the European Union’s Horizon Europe research and innovation programme (grant agreement no. 101082304 BlueRemediomics). We also thank the EMBL IT Services staff for managing and provision of access to the HPC resources.

## Supplementary Materials

Supplementary content file

Supplementary figures S1-S4

Supplementary tables S1-S10

Supplementary html files S1-S12

