## Supplementary figures for "GEMsembler: cross-tool structural comparison and ensemble modeling improve metabolic model performance"

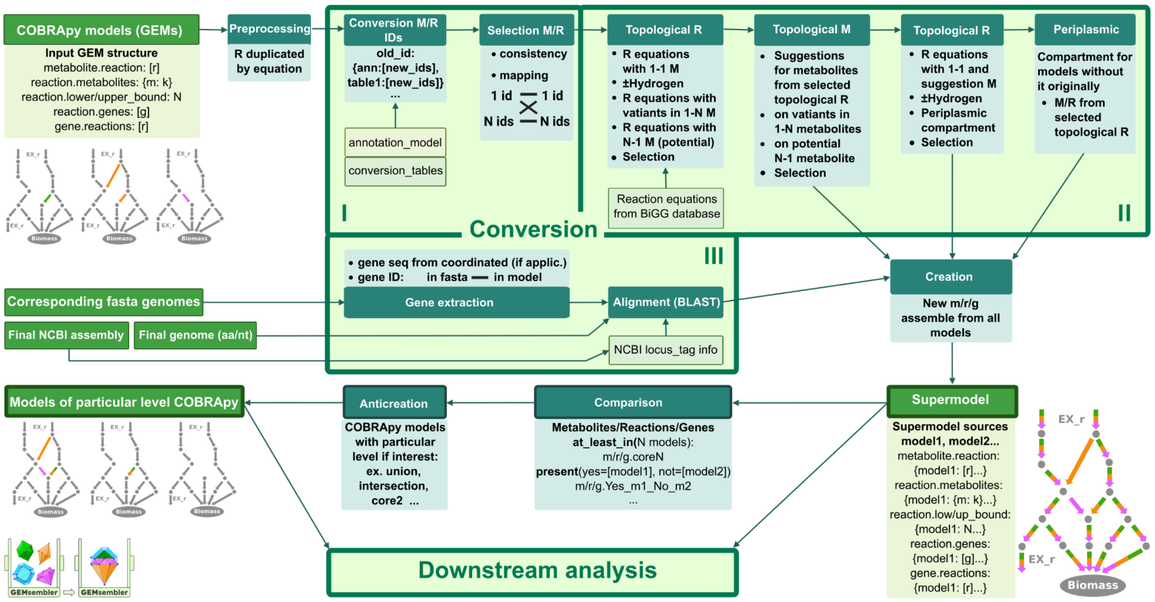


Figure S1. Schematic GEMsembler workflow.

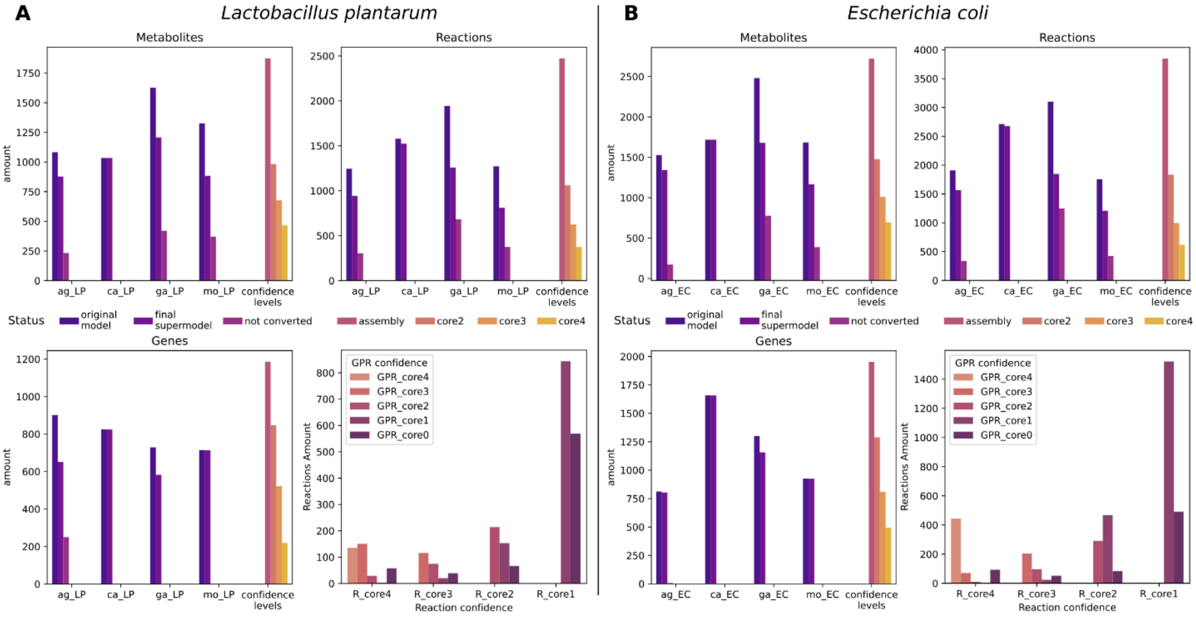


Figure S2. General characteristics of *L. plantarum* (A) and *E. coli* (B) models in terms of model agreement for metabolites, reactions, genes and GPRs. Ag: AGORA, ca: CarveMe, ga: gapseq, mo: modelSEED. CoreX corresponds to the agreement of X models.

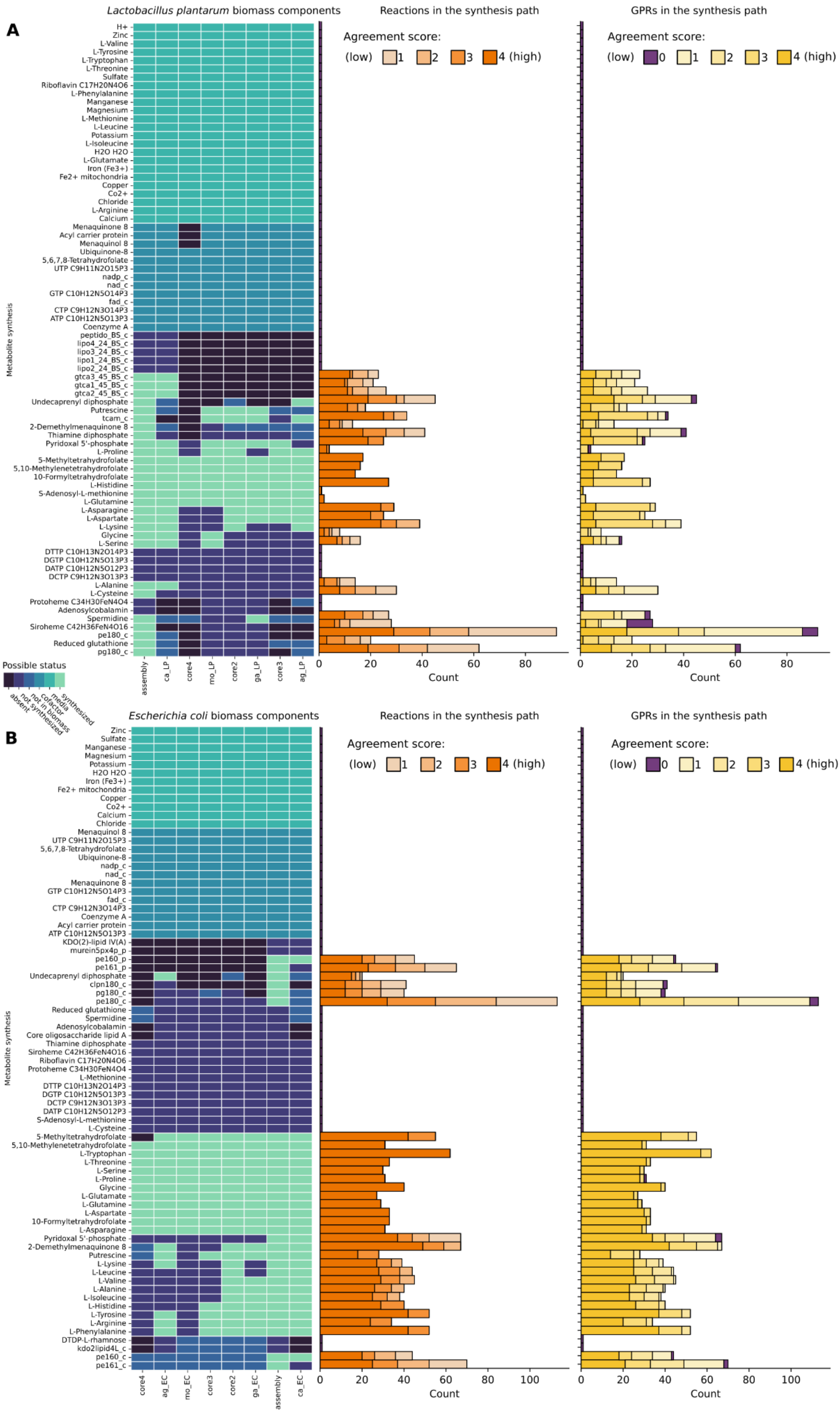


Figure S3. Biomass components production confidence for *L. plantarum* (A) and *E. coli* (B) models.

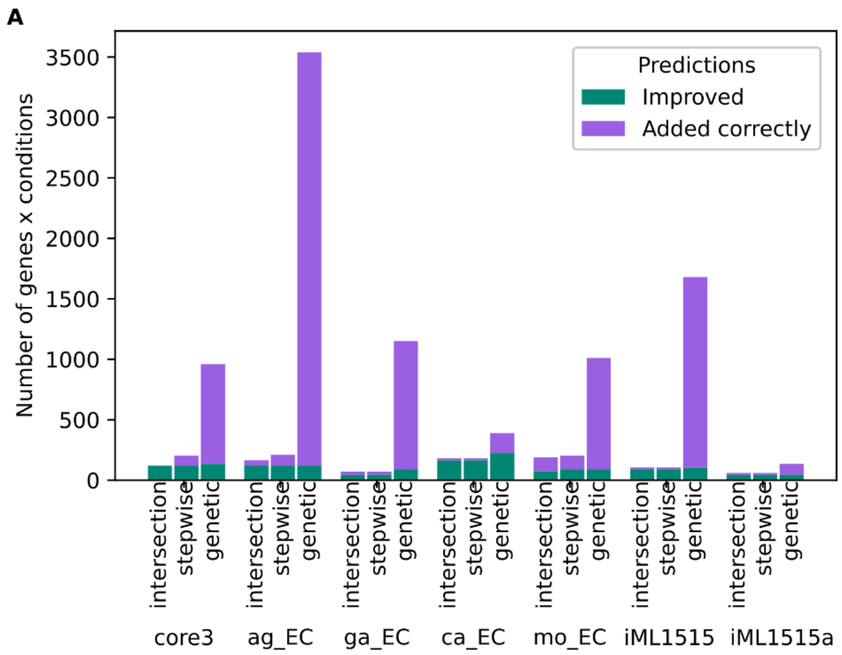


Figure S4. Number of [genes x condition] pairs (for carbon sources) for which the essentiality prediction improved using the models modified with either SA or GA or both of them; and [genes x condition] pairs (for carbon sources) with correct essentiality predictions for which the genes were newly added to each model by these algorithms.
