## Supplementary material for "GEMsembler: cross-tool structural comparison and ensemble modeling improve metabolic model performance": Content readme for supplementary tables and files on zenodo

Supplementary Tables
EC - *Escherichia coli*, LP - *Lactiplantibacillus plantarum.*
./Output/Supplementary_tables/

1. Supplementary Table 1. General statistics on the numbers in categories from different metabolic models for
    a. Metabolites LP (figS2a)
    b. Reactions LP (figS2a)
    c. Genes LP (figS2a)
    d. Reaction-GPR pairs LP (figS2a)
    e. Metabolites EC (figS2b)
    f. Reactions EC (figS2b)
    g. Genes EC (figS2b)
    h. Reaction-GPR pairs EC (figS2b)
2. Supplementary Table 2. Topology-determined (MetQuest) production of central carbon metabolites and their confidence
    a. Production status table LP (fig2a).
    b. Production status table EC (fig2b)
    c. Path confidence LP (fig2a)
    d. Path confidence EC (fig2b)
    e. The most confident production LP
    f. The most confident production EC
    g. Production statistics LP
    h. Production statistics EC
3. Supplementary Table 3. Predefined central carbon metabolism pathways and their confidence
    a. Glycolysis LP
    b. Glycolysis EC
    c. Pentose phosphate pathway LP
    d. Pentose phosphate pathway EC
    e. Tricarboxylic acid cycle (TCA) LP
    f. Tricarboxylic acid cycle (TCA) EC
4. Supplementary Table 4. Topology-determined (MetQuest) production of biomass components and their confidence
    a. Production status table LP (figS3a)

    b. Production status table EC (figS3b)
    c. Path confidence LP (figS3a)
    d. Path confidence EC (figS3b)
    e. The most confident production LP
    f. The most confident production EC
    g. Production statistics LP
    h. Production statistics EC
5. Supplementary Table 5. Biomass reaction composition, its confidence and decision on inclusion in the final biomass reaction
    a. Biomass composition and decision LP
    b. Biomass composition and decision EC
6. Supplementary Table 6. Summary of topology-based confidence analysis and identification of uncertainties
    a. Classification for metabolites of interest and their biosynthesis (fig2C)
    b. The most unconfident reactions (fig2D)
    c. Example of succinate biosynthesis LP (fig2E)
    d. Example of valine biosynthesis EC (fig2F)
7. Supplementary Table 7. Media composition
    a. PMM5 minimal media LP; CDPM media LP; M9 minimal media EC
8. Supplementary Table 8. Flux-determined (FBA/pFBA) production of biomass components and their confidence
    a. Production status table final biomass LP (fig3a)
    b. Production status table final biomass EC (fig3b)
    c. Example of interactive map for thiamine diphosphate curation (fig 3c)
    d. Production status table LP
    e. Production status table EC
    f. Production status table mixed LP (with non-converted features from the original model added to the converted model)
    g. Production status table mixed EC (with non-converted features from the original model added to the converted model)
9. Supplementary Table 9. LP models performance and comparison with gold-standard model
    a. Similarity table for reactions and the corresponding genes (fig4a)
    b. Number of reactions and genes from different models (fig4a)
    c. Auxotrophy growth status (fig4b) (0 - no growth, 1 - reduced growth, 2 - growth)
    d. Auxotrophy growth FBA simulations (flux value through the biomass reaction)
    e. Auxotrophy models performance (1 - match with the experimental data; 0 - mismatch)
10. Supplementary Table 10. EC models performance and comparison with gold-standard models
    a. Similarity table for reactions and the corresponding genes (fig5a)
    b. Number of reactions and genes from different models (fig5a)
    c. Models growth on different carbon sources
    d. AUCPR for each step in SA (stepwise procedure algorithm)

e. Number of generations intersected for GA (genetic algorithm) solution
f. Number of GPR changed by SA or GA (fig5d)
g. Number of genes changed by SA or GA (fig5e)
h. Status of gene essentiality predictions (fig5f)
i. Gene prediction improvement (fig5g)
j. Gene-condition pairs prediction improvement (figS4)
k. Models growth on different nitrogen sources

All production tables (in Tables S2, S4, S8) use the following numeric code for production status:
5 - synthesised;
4 - in the media;
3 - cofactor;
2 - not in the biomass;
1- not synthesised;
0 - not in the model.

Supplementary Files (html maps)
Supplementary files contain interactive pathway maps generated by GEMsembler that were used to produce the results and are referred to in the text. EC - *Escherichia coli*, LP - *Lactiplantibacillus plantarum.*
./Output/Supplementary_files/

1. Glycolysis LP
2. Glycolysis EC
3. Pentose phosphate pathway LP
4. Pentose phosphate pathway EC
5. Tricarboxylic acid cycle (TCA) LP
6. TCA EC
7. Succinate biosynthesis LP (Fig. 2E)
8. Valine biosynthesis EC (Fig. 2F)
9. Thiamine diphosphate pFBA pathway LP (Fig. 3C)
10. Pyridoxal 5'-phosphate biosynthesis with pyridoxamine from published iLP728 _LP model
11. Pyridoxal 5'-phosphate biosynthesis without pyridoxamine from GEMsembler-curated core3 LP model
12. Glutamate biosynthesis with pyridoxamine from published iLP728 LP model
